## Supplementary data for "MAGinator enables strain-level quantification of *de novo* MAGs"

### **Supplementary Data:** MAGinator enables strain-level quantification of *de novo* MAGs

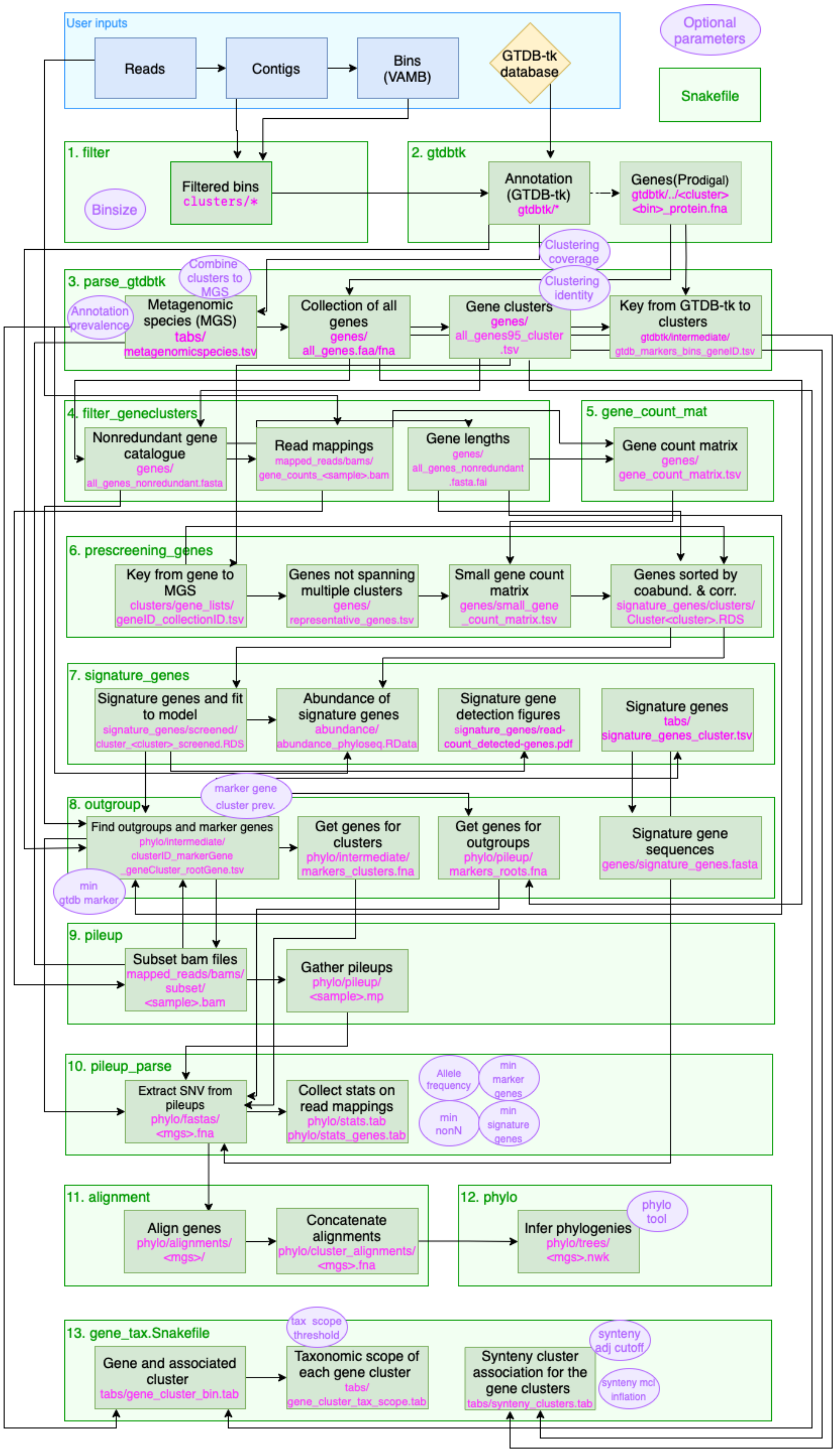
User inputs
Reads
Contigs
Bins (VAMB)
GTDB-tk database
Optional parameters
Snakefile
1. filter
Binsize
Filtered bins clusters/*
2. gtdbtk
Annotation (GTDB-tk) gtdbtk/*
Genes(Prodigal) gtdbtk/../<cluster> <bin>_protein.fna
Clustering coverage
Clustering identity
3. parse_gtdbtk
Combine clusters to MGS
Annotation prevalence
Metagenomic species (MGS) tabs/ metagenomicspecies.tsv
Collection of all genes genes/ all_genes.faa/fna
Gene clusters genes/ all_genes95_cluster .tsv
Key from GTDB-tk to clusters gtdbtk/intermediate/ gtdb_markers_bins_geneID.tsv
4. filter_geneclusters
Nonredundant gene catalogue genes/ all_genes_nonredundant.fasta
Read mappings mapped_reads/bams/ gene_counts_<sample>.bam
Gene lengths genes/ all_genes_nonredundant .fasta.fai
5. gene_count_mat
Gene count matrix genes/ gene_count_matrix.tsv
6. prescreening_genes
Key from gene to MGS clusters/gene_lists/ geneID_collectionID.tsv
Genes not spanning multiple clusters genes/ representative_genes.tsv
Small gene count matrix genes/small_gene _count_matrix.tsv
Genes sorted by coabund. & corr. signature_genes/clusters/ Cluster<cluster>.RDS
7. signature_genes
Signature genes and fit to model signature_genes/screened/ cluster_<cluster>_screened.RDS
Abundance of signature genes abundance/ abundance_phyloseq.RData
Signature gene detection figures signature_genes/read-count_detected-genes.pdf
Signature genes tabs/ signature_genes_cluster.tsv
marker gene cluster prev.
8. outgroup
Find outgroups and marker genes phylo/intermediate/ clusterID_markerGene geneCluster_rootGene.tsv
min gtdb marker
Get genes for clusters phylo/intermediate/ markers_clusters.fna
Get genes for outgroups phylo/pileup/ markers_roots.fna
Signature gene sequences genes/signature_genes.fasta
9. pileup
Subset bam files mapped_reads/bams/ subset/ <sample>.bam
Gather pileups phylo/pileup/ <sample>.mp
10. pileup_parse
Extract SNV from pileups phylo/fastas/ <mgs>.fna
Collect stats on read mappings phylo/stats.tab phylo/stats_genes.tab
Allele frequency
min marker genes
min nonN
min signature genes
11. alignment
Align genes phylo/alignments/ <mgs>/
Concatenate alignments phylo/cluster_alignments/ <mgs>.fna
12. phylo
Infer phylogenies phylo/trees/ <mgs>.nwk
phylo tool
13. gene_tax.Snakefile
tax scope threshold
Gene and associated cluster tabs/gene_cluster_bin.tab
Taxonomic scope of each gene cluster tabs/ gene_cluster_tax_scope.tab
Synteny cluster association for the gene clusters tabs/synteny_clusters.tab
synteny adj cutoff
synteny mcl inflation

***Supplementary Figure 1:*** *MAGinator workflow.*

A light green box indicates a snakefile and the darker green box indicates a deliverable (directory or file). The purple circles indicate user configurable parameters. The arrow indicates data dependencies, where the flow of information from one file is used to create the file it points towards.

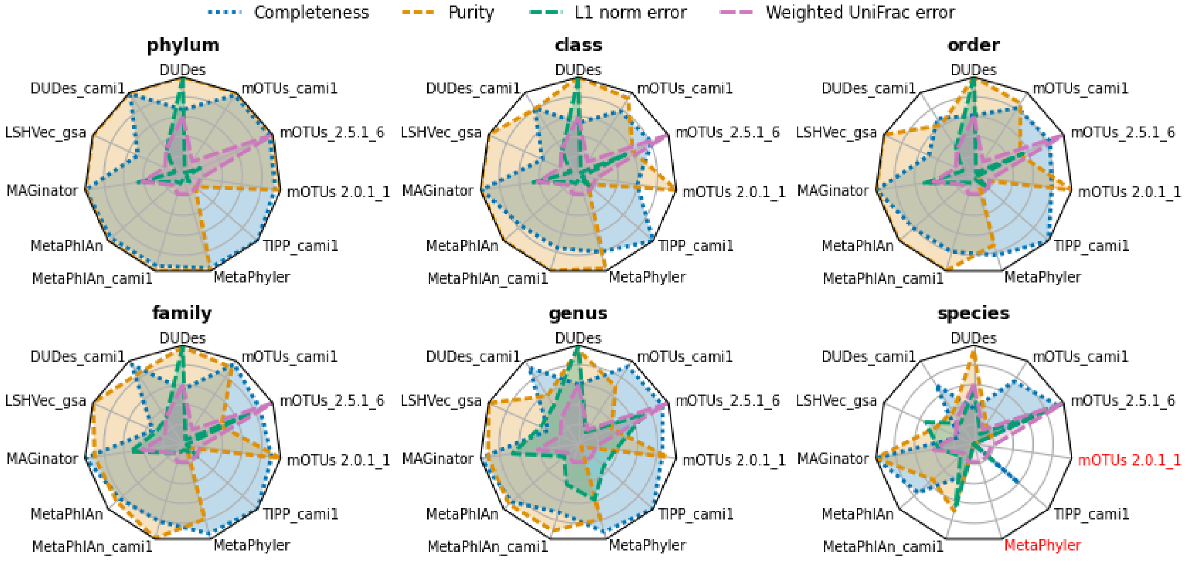

***Supplementary Figure 2****: Benchmark using OPAL*

Comparing taxonomic profiling results for the CAMI strain-madness data set. Metrics for the relative abundance of the profiles are calculated across samples for Completeness, Purity, L1 norm error and Weighted UniFrac error and are shown in a Spider-plot for the taxonomic ranks between phylum and species-level. The tools indicated with red means no data was available for that rank.

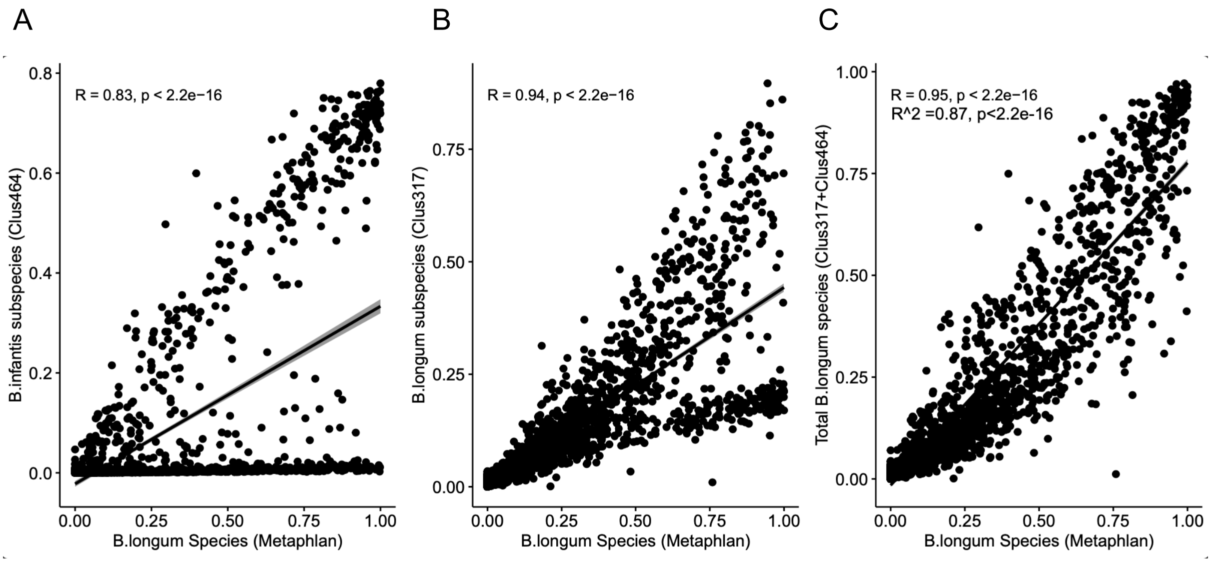

***Supplementary Figure 3:*** *Strain-resolved tracking of B. infantis metagenomes*

The relationship between the relative abundance for the *B. longum* species in the CHILD dataset found with MetaPhlAn and MAGinator's representative clusters for (A) *B. longum* (B) *B. infantis* (C) Both abundances added together. Each dot indicates a sample.

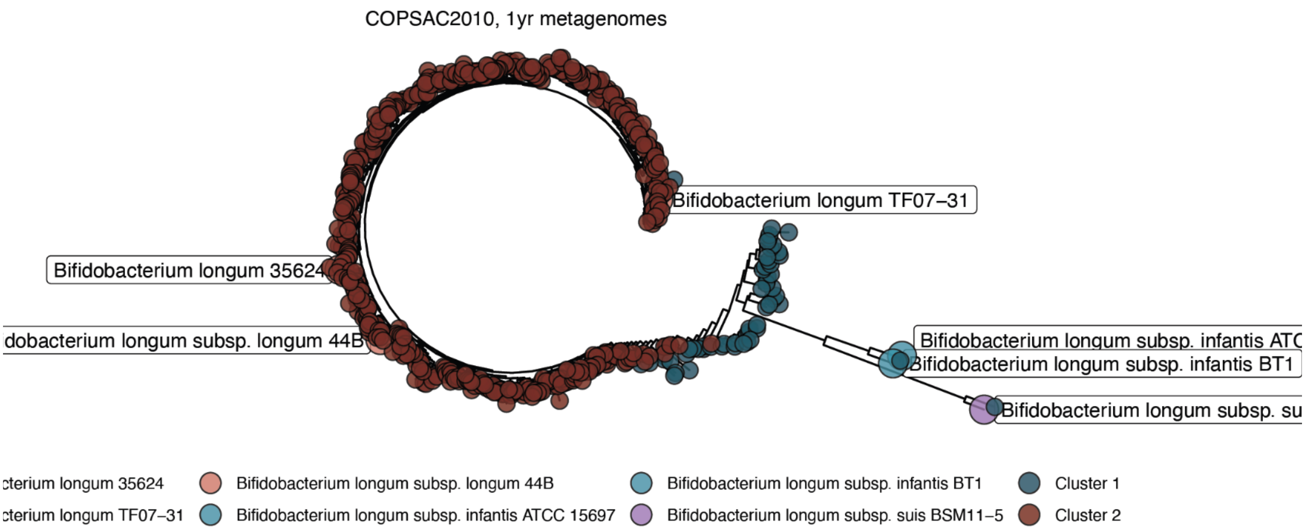

***Supplementary Figure 4****: Strain-level analysis of B. longum* in $COPSAC_{2010}$

StrainPhlAn phylogenetic tree of samples based on SNVs of B. longum markers, resulting in 2 clades. Dots represent samples with sufficient marker coverage as well as the 6 references. Cluster 1 indicates *B. infantis* and Cluster 2 indicates *B. longum.*

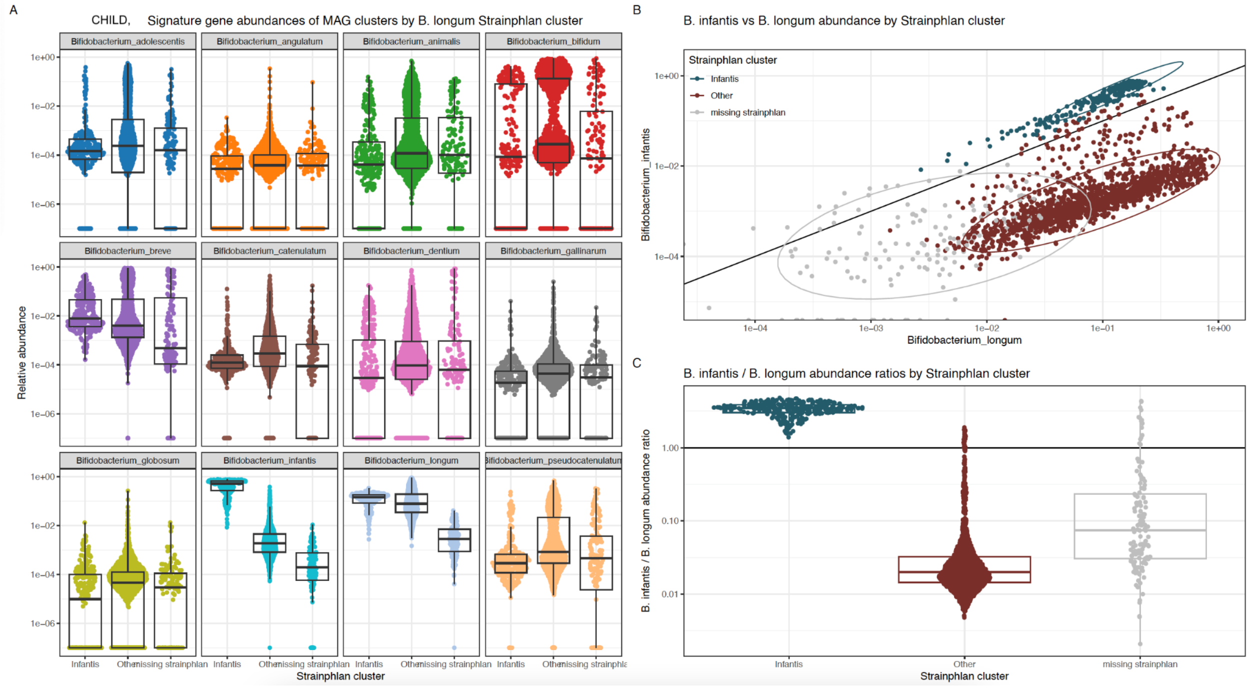

***Supplementary Figure 5:*** *Stratification of StrainPhlAn clusters using relative abundance of MAGinator clusters - CHILD cohort*

*Cluster 1 indicates B. infantis and Cluster 2 indicates B. longum.*

*(A) Relative abundance of StrainPhlAn clusters stratified by all Bifidobacterium clusters identified by MAGinator (B) Relative abundance of B. infantis and B. longum identified with MAGinator coloured by StrainPhlAn cluster. (C) The ratio of B. infantis to B. longum is displayed for the StrainPhlAn clusters.*

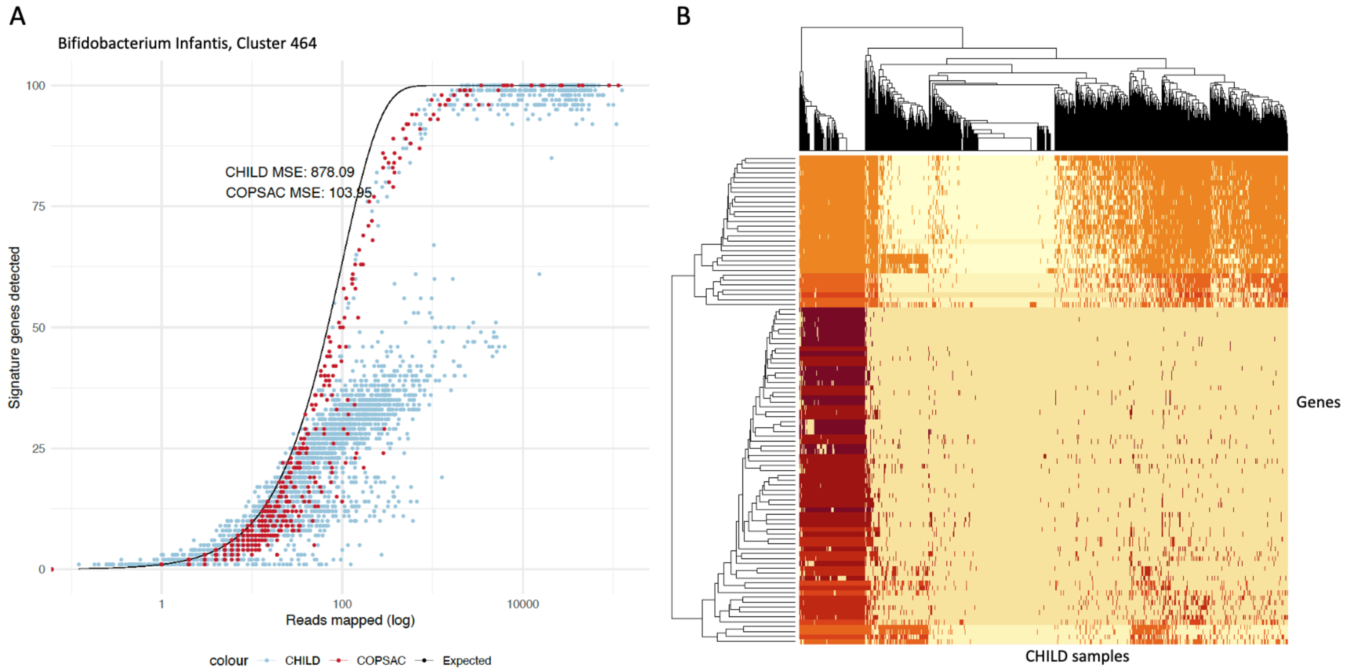

***Supplementary Figure 6:*** *Read mappings of B. infantis signature genes*

(A) The number of reads mapped to the signature genes (defined in $COPSAC_{2010}$) of the *B. infantis* cluster is presented with the number of signature genes detected. Each dot is a sample. The red colour indicates $COPSAC_{2010}$ samples, the blue colour indicates CHILD samples. The black line indicates the expected distribution[1]. (B) Heatmap of the read mappings of the *B. infantis* signature genes for the CHILD samples.

[1] Zachariasen, T. *et al.* Identification of representative species-specific genes for abundance measurements. *Bioinforma. Adv.* **3**, vbad060 (2023).

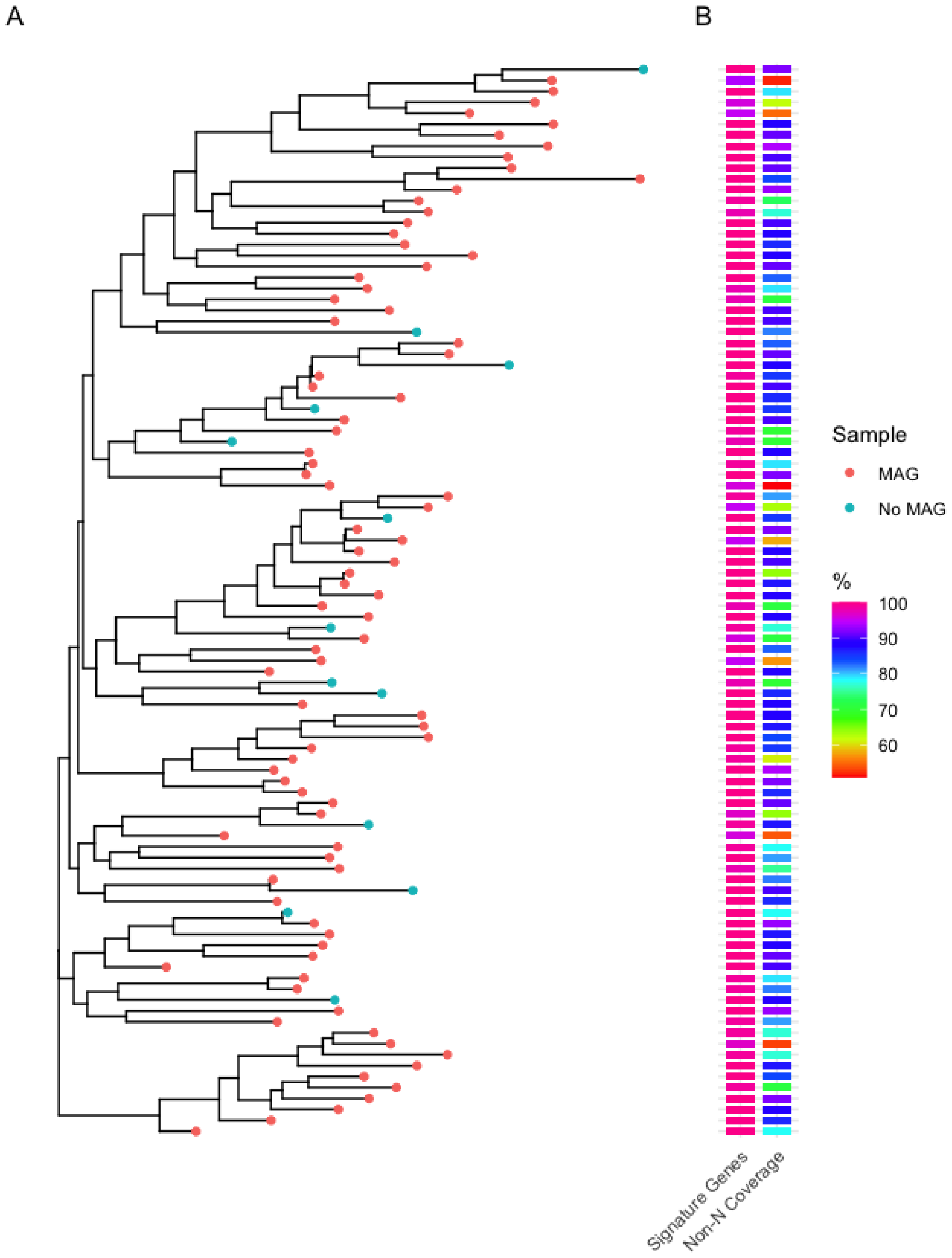

***Supplementary Figure 7**: SNV-level phylogenetic tree of a MAG cluster based on signature genes*

A) Phylogenetic tree constructed from readmappings to the signature genes of a MAG cluster annotated as *Faecalibacterium* sp900758465 from $COPSAC_{2010}$. Tip colour indicates if the sample has a MAG. B) Heatmap showing how many of the 100 signature genes that are detected within the sample and the fraction of bases that are Non-N in the alignment of the reads to the signature gene sequence in each sample (%).

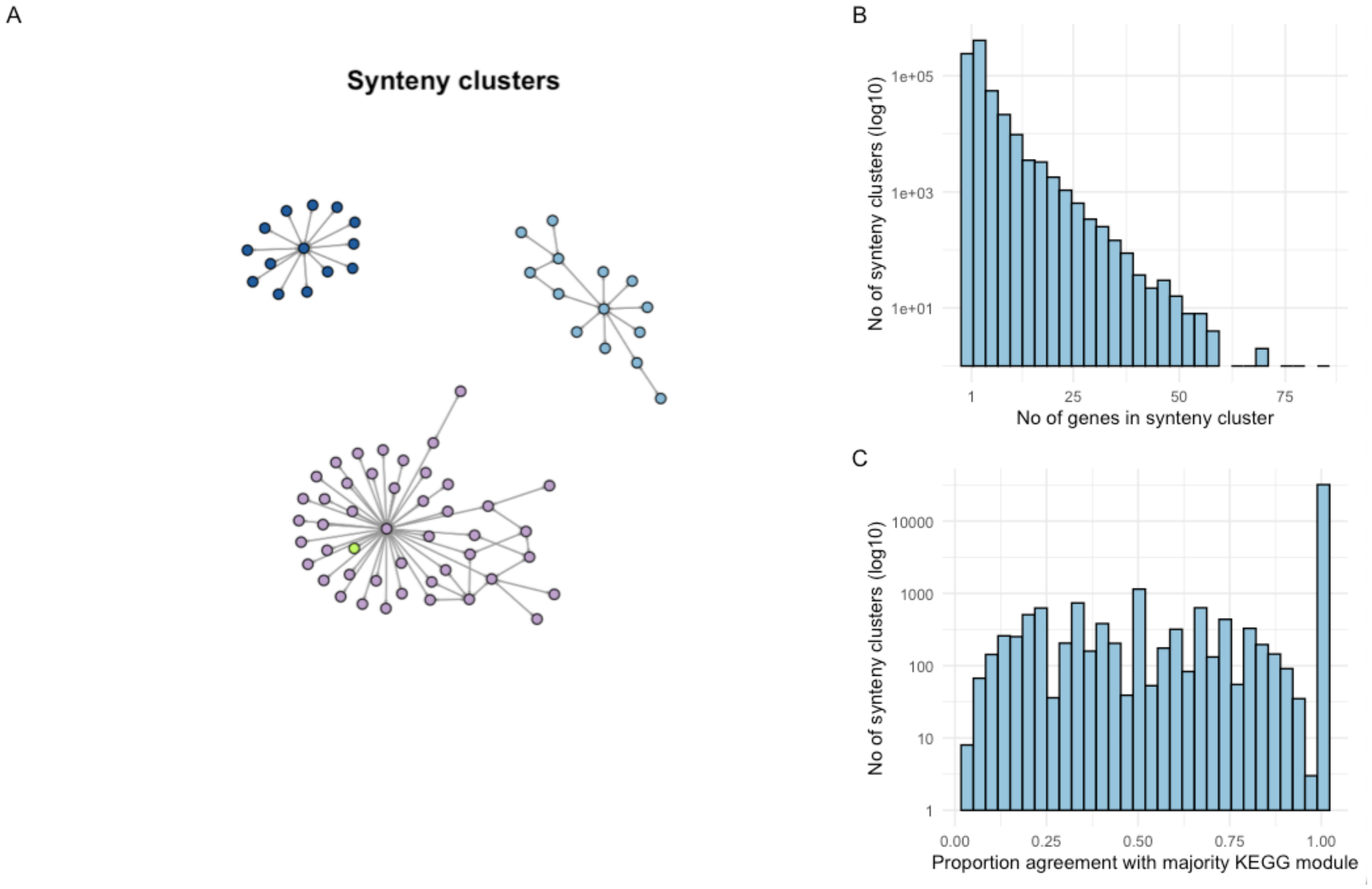

***Supplementary Figure 8:*** *Synteny clusters and functional annotation of COPSAC$_{2010}$*

A) The graph network of 3 synteny clusters are shown. The colours represent KEGG modules (green indicates no KEGG module annotation). B) The distribution of synteny cluster size. C) The proportion of genes in the synteny cluster in agreement with the most common KEGG module in the cluster. Only synteny clusters with 5 or more genes are included.

***Supplementary Table 1**: Output generated by MAGinator*

| Directory | Content |
|---|---|
| abundance | abundance_phyloseq.RData - Phyloseq object for R, with abundance and taxonomic data |
| clusters | .fa - Fasta files with nucleotide sequence of bins |
| genes | all_genes.faa - Amino acid sequences of all ORFs<br>all_genes.fna - Nucleotide sequences of all ORFs<br>all_genes_nonredundant.fasta - Nucleotide sequences of gene cluster representatives<br>all_genes_cluster.tsv - Gene clusters<br>matrix/gene_count_matrix.tsv - Read count for each gene cluster for each sample<br>synteny/ - Intermediate files for synteny clustering of gene clusters |
| gtdbtk | GTDB-tk taxonomic annotation for each MAG cluster |
| logs | Log files |
| mapped_ reads | bams/ - Bam files for mapping reads to gene clusters |
| phylo | alignments/ - Alignments for each signature gene<br>cluster_alignments/ - Concatenated alignments for each MAG cluster<br>pileup/ - SNV information for each MAG cluster and each sample<br>trees/ - Phylogenetic trees for each MAG cluster<br>stats.tab - Mapping information such as non-N fraction, number of signature genes and marker genes, read depth, and number of bases not reaching allele frequency cutoff<br>stats_genes.tab - Same as above but the information is split per gene |
| signature_g enes | R data files with signature gene optimization<br>read-count_detected-genes.pdf - Figure for each MAG cluster displaying number of identified SG's in each sample along with the number of reads mapped. |
| tabs | gene_cluster_bins.tab - Table listing which bins each gene cluster was found in<br>gene_cluster_tax_scope.tab - Table listing the taxonomic scope of each gene cluster<br>metagenomicspecies.tab - Table listing which, if any, clusters where merged in MAG cluster and the taxonomy of those<br>signature_genes_cluster.tsv - Table with the signature genes for each MAG cluster<br>synteny_clusters.tab - Table listing the synteny cluster association for the gene clusters. Gene clusters from the same synteny cluster are genomically adjacent.<br>tax_matrix.tsv - Table with taxonomy information for MAG cluster |

***Supplementary Table 2:*** *OPAL benchmark. Average completeness (%) across taxonomic ranks.*

The mean of the tools are indicated with bold.

| | superkingdom | phylum | class | order | family | genus | species | strain |
|---|---|---|---|---|---|---|---|---|
| **Mean across tools** | **100** | **90.1** | **75.1** | **79.0** | **83.3** | **82.7** | **45.6** | **0.7** |
| DUDes | 100 | 67 | 53.6 | 61.3 | 55.1 | 60.2 | 32.4 | 0 |
| DUDes cami1 | 100 | 98.8 | 81.6 | 68.8 | 95.7 | 91.8 | 63.4 | 0 |
| LSHVec gsa | 100 | 50 | 40 | 50 | 33.3 | 33.3 | 15 | 0 |
| MAGinator | 100 | 99.5 | 99 | 99.2 | 99.6 | 99.8 | 89.6 | 7.4 |
| MetaPhlAn | 100 | 96 | 76.8 | 80.7 | 84.3 | 89.2 | 67.1 | 0 |
| MetaPhlAn_cami1 | 100 | 94.8 | 75.8 | 79.8 | 81.8 | 71.6 | 29.4 | 0 |
| MetaPhyler | 100 | 97 | 80 | 83.3 | 94.2 | 92.2 | 0 | 0 |
| mOTUs 2.0.1_1 | 100 | 94.2 | 60 | 79.5 | 90.8 | 85.7 | 0 | 0 |
| mOTUs 2.5.1_6 | 100 | 97 | 81.6 | 84.7 | 85.9 | 93.6 | 84 | 0 |
| mOTUs cami1 | 100 | 97 | 78 | 81.7 | 96 | 92.9 | 67.4 | 0 |
| TIPP cami1 | 100 | 100 | 100 | 100 | 100 | 99.5 | 53.8 | 0 |

***Supplementary Table 3****: OPAL benchmark. Average purity (%) across taxonomic ranks.*

The mean of the tools are indicated with bold.

| | superkingdom | phylum | class | order | family | genus | species | strain |
|---|---|---|---|---|---|---|---|---|
| **Mean across tools** | **95.2** | **92.4** | **85.9** | **79.9** | **80.0** | **73.6** | **36.5** | **8.8** |
| DUDes | 100 | 100 | 100 | 100 | 98.1 | 95.3 | 84.9 | 0 |
| DUDes cami1 | 100 | 100 | 80.6 | 61.1 | 84.3 | 57.8 | 34.9 | 0 |
| LSHVec gsa | 100 | 100 | 100 | 100 | 100 | 100 | 37.5 | 0 |
| MAGinator | 100 | 100 | 100 | 100 | 90.3 | 92.4 | 90.1 | 96.7 |
| MetaPhlAn | 100 | 100 | 100 | 100 | 88.5 | 92.8 | 49.5 | 0 |
| MetaPhlAn cami1 | 97 | 100 | 100 | 100 | 99.3 | 91.1 | 63.8 | 0 |
| MetaPhyler | 100 | 100 | 97.4 | 72.7 | 78.9 | 79.2 | 0 | 0 |
| mOTUs 2.0.1_1 | 100 | 100 | 100 | 100 | 100 | 90.9 | 0 | 0 |
| mOTUs 2.5.1_6 | 100 | 100 | 60.6 | 51.5 | 43 | 37.3 | 17.8 | 0 |
| mOTUs cami1 | 100 | 100 | 93.2 | 87.4 | 91.7 | 69.1 | 21 | 0 |
| TIPP cami1 | 50 | 16.9 | 12.6 | 6.1 | 5.43 | 4.21 | 2.43 | 0 |
